# Dynamic network reconfigurations during task engagement following resting state in adolescent onset schizophrenia

**DOI:** 10.64898/2026.06.05.730513

**Authors:** Ouyang Bowei, Nicholas Theis, Jonathan Rubin, Konasale M. Prasad

**Affiliations:** Department of Psychiatry, University of Pittsburgh School of Medicine, Pittsburgh, PA; Department of Mathematics, Ken Dietrich School of Arts and Sciences, University of Pittsburgh, PA; Department of Bioengineering, University of Pittsburgh Swanson School of Engineering, Pittsburgh, PA; VA Pittsburgh Healthcare System, Pittsburgh, PA

**Author notes:** Corresponding author: 3811 O’Hara St, Pittsburgh, PA.

**Keywords:** Resting-state fMRI, Energy landscape analysis, Dynamical system, Brain state dynamics, Default mode network, Executive function, Maximum entropy model

## Abstract

Understanding how functional brain networks in resting state configurations reorganize to perform cognitive tasks is critical for uncovering the nature and mechanisms underlying network dysconnectivity in psychiatric disorders. We applied energy landscape analysis (ELA), a statistical physics-based computational approach, to functional MRI data from 23 adolescent-onset schizophrenia (AOS) and 44 healthy control (HC) subjects, acquired during rest followed by executive function task. ELA maps brain activity into distinct network states and quantifies how the brain transitions among them, capturing differences in stability of network states and transition complexity across conditions. AOS and HC showed markedly different condition-dependent patterns of brain state organization and dynamics. At rest, AOS exhibited reduced dynamical complexity compared to HC (7 vs. 14 stable states) that reversed during the task with more than 2-fold increase in accessible but rarely occupied brain states, while HC showed an opposite pattern. These results suggest that cognitive demands unmask latent fragmentation of the energy landscape, comprising a proliferation of accessible but rarely occupied states not apparent at rest, in AOS. State occupancy analysis revealed a small number of dominant states accounting for the majority of brain activity time, with AOS showing greater persistence in the fully-active DMN state during task performance compared to HC. These findings suggest that the rest-to-task transition features fundamentally different neural dynamics in AOS compared to HC. Combined analysis of resting fMRI and task-induced brain dynamics revealed neural factors that may contribute to cognitive dysfunction and psychiatric symptoms in schizophrenia, with important implications for development of biomarkers and treatment targets.

## 1. Introduction

Resting-state functional connectivity (rsFC) measures how spontaneous blood oxygenation level dependent (BOLD) signal fluctuations in spatially distinct brain regions are temporally correlated when a person is not performing any explicit task. The BOLD signal fluctuations are not random but are organized into several resting state networks (RSNs) that interact with each other and show distinct features in health and disease states. One such interaction that is well-studied is the suppression of activity in default mode network (DMN) regions when an individual engages in an attention-demanding task. In schizophrenia, the DMN regions often show hyperactivation with reduced task-related suppression, which has been associated with cognitive impairments (Menon, 2023; Whitfield-Gabrieli & Ford, 2012). Since DMN activity is typically quantified using pairwise correlation of BOLD signals across DMN regions averaged over the entire duration of acquisition, traditional connectivity approaches may obscure dynamic changes in regional activation/deactivation over shorter timescales such as at each repetition time (TR) of fMRI acquisition. Similarly, temporal averaging over a task block does not reveal whether regions consistently alter their activity during the entire duration of data acquisition or task blocks, merely do so on average across variable timepoints, or dynamically change while performing the task. While traditional dynamic functional connectivity approaches can capture time-varying changes by computing connectivity within sliding windows, these methods face methodological challenges including dependence on the choices of window length and sliding interval, which can limit temporal resolution, and the possibility that high within-window correlations may arise when two regions are simultaneously inactive, reflecting shared quiescence rather than functional coupling, and therefore may be less functionally meaningful.

Energy landscape analysis (ELA) offers an integrated and comprehensive complementary approach by directly modeling the changes in brain states (described below) at every TR of fMRI data without requiring arbitrary window definitions. ELA uses statistical physics principles, which model the behavior of a large collection of microstates to understand the macroscopic behavior at the systems level (Brush, 1967; E. Ising, 1924; T. Ising, Folk, Kenna, Berche, & Holovatch, 2017). ELA characterizes network dynamics by modeling neural activity as the path a brain takes through different activity patterns over time, where the fMRI signal at each moment in time defines the brain’s current “position” on this landscape (Ishida et al., 2024). This approach uses “energy” (a statistical analog of Gibbs free energy, a measure of how probable a given brain activity pattern is, not physical or metabolic energy) to describe brain states (distinct patterns of brain activity, each representing a unique combination of regions being active or inactive at a given moment). The energy term is inversely related to the occurrence probability of network configurations such that higher energy is associated with network configurations of lower probability and vice versa. According to this conceptualization, the brain network system naturally gravitates toward low-energy stable (self-sustaining) states (local minima or basins of attraction) but can transition among states due to perturbations by noise, inputs from other regions, external stimuli, or spontaneous fluctuations. ELA maps the full landscape of brain states including both stable (patterns of activity the brain repeatedly returns to, i.e., low energy states) and unstable or transient states (patterns of activity that the brain passes through briefly while shifting between stable configurations). “Transition dynamics” refers to how the network moves between these states: which states it switches between, along what pathways states evolve, and the dwell times or duration of time spent in each state. Just as a ball rolling across a hilly landscape will tend to settle in valleys but can roll from one valley to another when pushed, the brain activity tends to settle into stable states but can shift between them over time. ELA quantifies three critical features of brain dynamics (Ezaki, Watanabe, Ohzeki, & Masuda, 2017): (1) the number and energy depth of stable states (local minima), (2) state occupancy and dwell times, and (3) transition rates and pathways between states.

While these measures have been examined in autism spectrum (Watanabe & Rees, 2017), showing altered dynamics and reduced flexibility in state transitions, their application to schizophrenia has remained limited, particularly in adolescent-onset schizophrenia (AOS) and in rest-to-task-dependent switches. Indeed, recent fMRI studies in schizophrenia have revealed aberrant neural dynamics beyond static connectivity alterations (Friston, Harrison, & Penny, 2003; Rolls & Deco, 2011), suggesting that cognitive impairments may arise from disrupted temporal organization of brain states rather than simply altered connection strengths (Deco, Tononi, Boly, & Kringelbach, 2015; Tang et al., 2008). However, most energy landscape studies have focused on either resting-state (Ishida et al., 2024) or task fMRI data (Watanabe, Masuda, Megumi, Kanai, & Rees, 2014) separately, leaving unclear whether patients show condition-specific patterns that differ between rest and cognitive engagement compared to controls.

In the current study, we employed a data-driven ELA approach, with model parameters inferred directly from fMRI data, to examine brain state dynamics in AOS and age-matched healthy controls (HC) at rest and during performance of the Penn Conditional Exclusion Test (PCET), an executive function task (Gur, Ragland, Moberg, Bilker, et al., 2001; Gur, Ragland, Moberg, Turner, et al., 2001). We previously reported using the same dataset examining the executive function network that AOS patients showed higher energy levels with low basin volumes, that AOS trajectories traversed higher energy states while controls’ trajectories remained in lower energy domains, and that higher energy values correlated with reduced performance on PCET and greater severity of psychopathology (Theis et al., 2025). We focused on AOS because AOS is associated with especially prominent developmental and premorbid abnormalities with more severe cognitive impairments (Frangou, 2010; Holtmaat & Svoboda, 2009; Kester et al., 2006; Rapoport & Gogtay, 2011; Thaden et al., 2006), particularly in working memory (Brickman et al., 2004; Karatekin, Bingham, & White, 2009; Karatekin, White, & Bingham, 2008; White, Mous, & Karatekin, 2013) and executive functions (Frangou, 2010; Kester et al., 2006; Rapoport et al., 1997; Thaden et al., 2006) with poor long-term outcomes (Frangou, 2010; Kumra & Charles Schulz, 2008; Kumra, Shaw, Merka, Nakayama, & Augustin, 2001; Rapoport et al., 1997). Hence, AOS is proposed as a severe form of schizophrenia with potentially stronger neurodevelopmental underpinnings (Kumra & Charles Schulz, 2008; Musket et al., 2020). Despite this evidence, the neurobiology of AOS is under-investigated (Theis et al., 2025). In this study, our goals were to perform ELA using fMRI data for the DMN during rest and to describe the DMN landscape when subjects shift to task performance. During task epochs, we expected AOS to show higher energy compared to controls (Theis et al., 2025; Whitfield-Gabrieli & Ford, 2012). We also expected both AOS and HC to show lower energy during rest than during task performance due to higher energy associated with cognitive performance demands. While we observed the former, to our surprise, we found that the task-based reduction of energy was present in controls yet reversed in AOS, suggesting challenges in maintaining task engagement among AOS.

## 2. Material and methods

### (a) Participants and data acquisition

We enrolled AOS from inpatient and outpatient clinics at the University of Pittsburgh Medical Center, Pittsburgh, and age-matched HC from the community. AOS subjects had the onset of first psychotic symptoms after puberty but before reaching 19 years of age. Puberty was defined as scoring ≥2 on the Peterson Pubertal Developmental Scale (Petersen, Crockett, Richards, & Boxer, 1988). Participants diagnosed with intellectual disability according to the DSM-IV, having history of substance use disorder in the last 3 months, head injury with significant loss of consciousness, tumors, encephalitis, or suffering neonatal asphyxia were excluded. Subjects were administered the Structured Clinical Interview for DSM-IV (SCID-IV) and selected items on the Kiddie-Schedules for Assessment of Depression and Schizophrenia (K-SADS) (Kaufman, Birmaher, Brent, Ryan, & Rao, 2000). Consensus diagnosis by experienced clinical investigators was made after reviewing all available clinical data including the charts. The University of Pittsburgh Institutional Review Board approved the study. Parents or guardians provided informed consent for participants under 18 years of age, who also provided their assent; participants aged 18 or older provided consent directly. All were given a complete description of the study, including its risks and benefits. More details are in our prior publication (Theis et al., 2025)

Imaging data were obtained on a 7 Tesla whole-body scanner. T_1_-weighted MP2RAGE scans were acquired in the axial plane: 348 slices, 0.55mm thickness, TE=2.54ms, TR=6000ms, in-plane voxel matrix size of 390×390, at 0.55mm isotropic voxels resolution. Two functional MRI (fMRI) sequences were acquired: one in resting conditions and another during performance of PCET for executive function. Both were functional echo-planar images acquired in the axial plane with isotropic voxel size of 1.5mm for PCET and 2mm for rest. For the resting state fMRI, 60 slices of 2mm thickness, TE=20ms, TR=1000ms, and an in-plane resolution of 110 × 110 were collected, totaling 480 volumes (8 minutes, 16 seconds). For PCET, 86 slices of 1.5 mm thickness, with TE=20ms, TR=3000ms, and in-plane resolution of 148 × 148 were collected, totaling 142 volumes (7 minutes, 33 seconds). Field maps were acquired for magnetic susceptibility distortion correction with the same acquisition parameters as each fMRI in terms of matrix size and slab number, but with a longer TE (36.2ms) and TR (6000ms).

### (b) Data preprocessing and region-of-interest (ROI) extraction

FreeSurfer (Fischl, 2012) was used to preprocess and parcellate the T_1_w images. The 360-node Glasser parcellation was applied (Glasser et al., 2016) to define cortical ROIs, and we used the standard FreeSurfer cortical and subcortical parcellation to bring the total to 383 ROIs. All pre-processing steps for the fMRI data were performed using fMRIPrep (Esteban et al., 2019), which included correction for motion, slice timing, bias, gradient distortion, and brain extraction, before registering to a standard space (MNI 2mm) using the Human Connectome Project approach (Glasser et al., 2013).

Image registration of T_1_w volumes to fMRI space was performed, and the transformation was then applied to the atlas using nearest-neighbor interpolation, yielding ROI labels in native fMRI space.For each brain region (parcel) in the registered atlas image, the voxel-wise BOLD signal of the preprocessed fMRI was calculated at each timepoint. This yielded a multivariate time series matrix for each subject and each fMRI acquisition representing activity of brain regions over time. These time series were then *z*-scored per subject per brain region (node) and subsequently binarized using a threshold of zero; regions showing BOLD signal intensity >0 were coded as “on/active” or 1 and ≤0 as “off/inactive” or 0. Finally, subsets of HCP/Glasser atlas-defined regions were selected for the DMN (Sandhu et al., 2021) for subsequent analysis.

We selected 10 (out of 34) left-hemisphere ROIs based on their high on/off switching rates during the scanning sessions: retrosplenial cortex (L_RSC), posterior cingulate regions (L_POS1, L_POS2), ventral posterior cingulate areas (L_v23ab, L_31pv), dorsal posterior cingulate areas (L_d23ab), anterior cingulate-medial prefrontal cortex (L_a24, L_p32, and L_10r), and temporoparietooccipital junction (L_TPOJ3). The switching rate was calculated for all 34 ROIs from the binarized time series, where switching was defined as change in states across two consecutive time steps (i.e, 0 to 1 or 1 to 0). The switch rate was calculated for every ROI as the ratio of the total number of switches to the total number of TRs. The ROIs were sorted in descending order (highest number of switches to the lowest) and the top 10 ROIs showing the highest number of switches were selected as the most dynamic ROIs that may be particularly informative for ELA. These DMN regions have also been associated with cognitive control (Cole, Bassett, Power, Braver, & Petersen, 2014). We supplemented this approach with 500 permutation tests by randomly selecting 10 ROIs out of a total of 34 to ensure that the results are not due to a bias in the node selection.

### (c) Pairwise maximum entropy model

We first specified a brain network of 10 ROIs and obtained BOLD signals from these regions, resulting in a multivariate time series of BOLD signals. Next, we normalized the fMRI signals by calculating a *z*-score at each time point for each ROI and then binarized these signals relative to a threshold at 0, resulting in a time-dependent binary sequence {σ_i_(1), …, σ_i_(*t*_*max*_)} where *t*_*max*_ denotes the total duration of the data sample, σ_i_(*t*) = 1 indicates that the *i*^th^ ROI is active at time *t*, and σ_i_(*t*) = 0 indicates that the ROI is inactive. The activity pattern of the entire network at time *t* is given by an N-dimensional vector ***σ*** ≡ (σ_1_, …, σ*N*) ∈ {0, 1}^N^. Note that there are 2^*N*^ possible state patterns in total, each of which could in theory be present at each time *t*. We next calculated the relative frequency with which each activity pattern is visited, *P*_*empirical*_(σ). To estimate the model-based probability distribution, we fitted the empirical frequency distribution with the Boltzmann distribution given by:

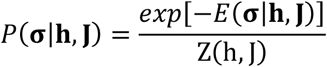

where 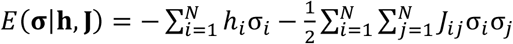 is the energy term, 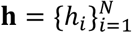 represents the external field parameters, i.e. regional BOLD signal level, and 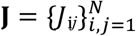 represents the pairwise interaction parameters. We assume *J*_*ij*_ = *J*_*ji*_ (symmetric coupling) and *J*_jj_ = 0 (no self-coupling). The partition function was defined as:

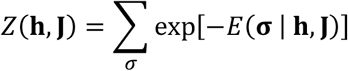

which normalizes the probability distribution. The principle of maximum entropy imposes that we select *h* and *J* such that the model reproduces the empirical first and second moments:

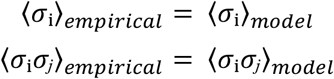

where ⟨·⟩_*empirical*_ and ⟨·⟩_*model*_ represent expectations with respect to the empirical and model distribution, respectively; with these choices, we refer to this model for state probabilities as the maximum entropy model (MEM).

### (d) Parameter estimation

We estimated the MEM parameters using an improved two-stage optimization approach that enhances convergence speed and stability compared to standard methods.

#### Step 1: Pseudolikelihood Initialization

We first obtained initial parameter estimates using the pseudolikelihood method (Ezaki et al., 2017). The pseudolikelihood method provides a computationally tractable approximation by simplifying the calculation: instead of computing probabilities across all 2^*N*^ possible states simultaneously (which becomes prohibitively expensive as *N* increases), it approximates the likelihood by considering each node’s activation probability conditioned on its neighbors. This avoids the expensive summation over all 2^*N*^ states required for exact likelihood calculation. Parameters are updated iteratively using the equations:

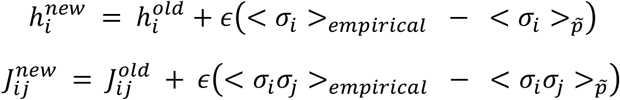

where ϵ is the learning rate (step size) that controls the magnitude of parameter updates and 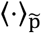 represents expectations with respect to the pseudolikelihood approximation.

#### Step 2: Refined Maximum-Likelihood Estimation

Using the pseudolikelihood estimates from Stage 1 as initial values, we performed refined optimization via the L-BFGS-B algorithm (Limited memory Broyden-Fletcher-Goldfarb-Shanno with Box constraints). Unlike the pseudolikelihood approximation, this stage computes the exact likelihood by evaluating probabilities across all possible states. This quasi-Newton method offers several advantages:

1. automatic step-size adaptation via line search,
2. approximation of the Hessian matrix for better convergence properties, and
3. memory-efficient implementation suitable for high-dimensional problems. We minimize the negative log-likelihood:

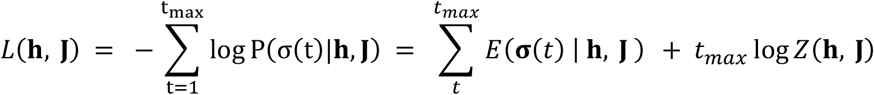

where *L*(**h**, J) represents the negative log-likelihood function that we seek to minimize and *t*_*max*_ is the total number of time points in the data.

The two-stage approach combines computational efficiency with accuracy: the pseudolikelihood initialization provides a computationally efficient starting point close to the optimal solution, while the L-BFGS-B refinement using exact likelihood ensures high-quality parameter estimates. This hybrid strategy substantially reduces the total number of expensive exact likelihood evaluations required compared to optimizing random initialization.

### (e) Disconnectivity graph and energy landscape construction

After fitting the pairwise MEM, we constructed disconnectivity graphs: hierarchical representations that depict the energy barriers separating different local minima, encapsulating both the stability of brain states and the difficulty of transitioning between them. Local energy minima are identified as activity patterns with smaller energy than that of all neighbors. The neighborhood of σ is defined as the set of *N* activity patterns with a Hamming distance of 1 from σ (i.e., differing from σ in the activation state of exactly one ROI). For a given pair of local minima, *a* and *b*, we consider paths connecting them, where a path is a sequence of activity patterns such that any two consecutive patterns are neighbors (Hamming distance=1). The energy barrier for the transition from *a* to *b* is calculated using a Dijkstra-like algorithm (Dijkstra, 1959) to find the minimum energy path.

From the disconnectivity graphs, two-dimensional energy landscape visualizations were constructed by first applying hierarchical clustering with Ward linkage to the disconnectivity matrix, which yielded a spatial arrangement of local minima in the 2D plane. Each local minimum refers to a numbered state, with inter-state distance reflecting the height of the energy barrier forming the boundary between the basins (see (f) below). Energy values were assigned at each state position and at the internal branch points of the clustering tree, then interpolated across a uniform 2D grid using inverse distance weighting to produce a continuous contour surface. The resulting surface was color-coded by energy level, with darker colors indicating lower energy.

### (f) Basin of attraction analysis

Each local minimum has a basin of attraction in the state space. We determined basin membership for each state pattern *s* using a gradient descent approach. Starting from any non-minimal state *s*, we iteratively transitioned to neighboring activity patterns with lower energy values, where each transition involved switching the activity status of a single ROI (changing one bit in the binary pattern). This process continued until the system reached a local energy minimum *m*, at which point we assigned state *s* to the basin of attraction of *m*. This procedure effectively mapped the state space into distinct basins, each corresponding to one of the identified local minima.

### (g) Model validation and statistical analysis

ELA was performed at the group level by concatenating binarized time series across all subjects within each group and condition prior to model fitting. This approach was necessitated by the dimensionality of the state space: with N=10 ROIs, there were 2^10^=1,024 possible binary configurations. Group-level concatenation is consistent with established practice in the ELA literature (Ezaki et al., 2017; Ishida et al., 2024; Watanabe & Rees, 2017) and reflects the collective dynamics of each group under each condition.

The quality of the MEM model fit was assessed using the correlation coefficient between empirical and fitted state probabilities. Specifically, the accuracy of the pairwise MEM was quantified using the Pearson correlation coefficient *r* between empirical and model-fitted state probabilities across all 2^10^ = 1,024 possible binary patterns. Group differences in the number of local minima, state frequencies, and basin sizes were evaluated. Direct transition matrices were computed to quantify the probability of transitioning between the basins of attraction of pairs of local minima.

## 3. Results

### (a) Model accuracy and validation

The pairwise maximum entropy models achieved excellent fits across all conditions and groups. For resting-state data, correlation coefficients between empirical and fitted probabilities were *r*=0.935 for patients and *r*=0.996 for controls (Supplemental Figures 1B, 1D). For PCET task data, correlations were *r*=0.969 for patients and *r*=0.988 for controls (Supplemental Figures 1A, 1C). The high correlations and tight clustering around the identity line confirm that the pairwise MEM accurately captured the observed brain state distributions.

### (b) Energy landscape dynamics in healthy controls

At rest, healthy controls exhibited a complex energy landscape with 14 local minima **(Figure 1A, B)**, with values of the dimensionless energy associated with the MEM (see Methods, part (c)) ranging from −8 to −15. The three most frequently occupied states (Figure 1D, left) were state 0 (∼41%), state 1023 (∼27%), and state 14 (∼19%). State 0 represented the all-off state, in which all 10 ROIs were simultaneously inactive (z-scored BOLD response ≤0), state 1023 showed complete DMN engagement with all 10 ROIs active, and state 14 had only the anterior cingulate and medial prefrontal regions active (L_a24, L_p32, and L_10r). The remaining states each occurred in less than 5% of measurements; 6 of the 14 states were predicted as minima by the model but not observed empirically. The 2D energy landscape **(Figure 1C)** reflected the overall complexity of the resting state dynamics, with the 14 local minima distributed across a broadly contoured surface organized around deep energy wells at states 0 and 1023 (Figure 1C). The transition matrix (Figure 1D, right) showed the dominant coupling to be between states 0 and 14, with high bidirectional transition counts, with additional transitions involving the less frequently observed state, i.e. 712, featured in transitions with state 0, and 271, featured in transitions with state 14. Since it was frequently observed but participated in very few transitions, we infer that state 1023 behaved as a nearly-absorbing state: the extremely low transition rate out of this state indicates that, once entered, the system tended to remain there for prolonged dwell times rather than cycling through other configurations.

**Figure 1:**
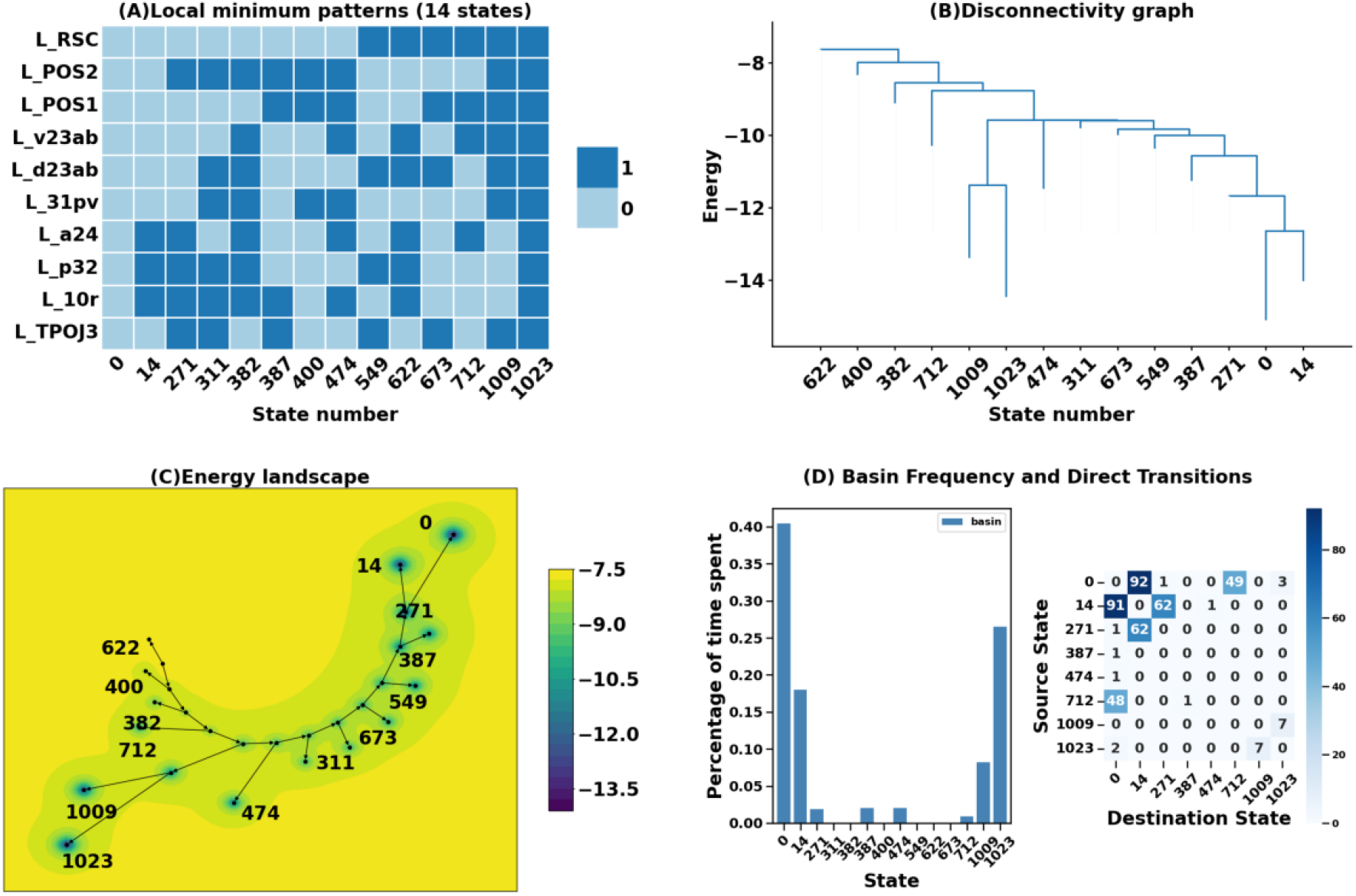
Energy landscape analysis in the resting state for control subjects. (A) Binary activation patterns for 14 local minima across 10 brain regions. (B) Hierarchical disconnectivity graph showing a more complex hierarchical organization with energy values ranging from -8 to -15. (C) Energy landscape visualization showing the spatial organization of local minima and energy gradients across state space. (D) *Left:* Basin frequency distribution showing the percentage of time spent in each basin, with states 0, 1023, and 14 most frequently occupied. States without bars evident represent model-predicted local minima with zero empirical occupancy. *Right:* Direct transition matrix quantifying transition probabilities between states.

During PCET, the energy landscape of healthy controls underwent a striking consolidation. The number of local minima dropped from 14 (with 8 observed empirically) at rest to 4 during task engagement **(Figure 2A, B)**, with energy values ranging from approximately −6 to −16. The dominant state was state 0 (∼47%), representing complete DMN suppression with all 10 ROIs simultaneously inactive. This all-off state was also the deepest minimum of the entire PCET landscape, reaching -16 **(Figure 2B)**. The second most frequent state was state 1023 (∼27%), representing complete DMN engagement with all 10 ROIs simultaneously active. State 14 (∼18%) showed sparse activation with only 3 of 10 ROIs active, exclusively the anterior cingulate and medial prefrontal regions (L_a24, L_p32, and L_10r), while state 1009 (∼10%) showed broader activation with 7 of 10 ROIs active (L_RSC, L_POS2, L_POS1, L_v23ab, L_d23ab, L_31pv, and L_TPOJ3). The disconnectivity graph **(Figure 2B)** revealed two well-separated hierarchical clusters: states 0 and 14 on the left (merging at approximately −14), and states 1023 and 1009 on the right (merging at approximately −14), with the two clusters separating at approximately −6. Transition dynamics corroborated this separation, with high numbers of transitions between states 0 and 14 as well as between states 1023 and 1009, in both directions **(Figure 2D)**. The 2D landscape **(Figure 2C)** reflected this organization with 4 nodes arranged into two spatially separated clusters. The latter set of transitions contrast with the absorbing character of state 1023 in rest conditions, suggesting that exit out of state 1023 may be necessary for, or at least associated with, successful task performance.

**Figure 2:**
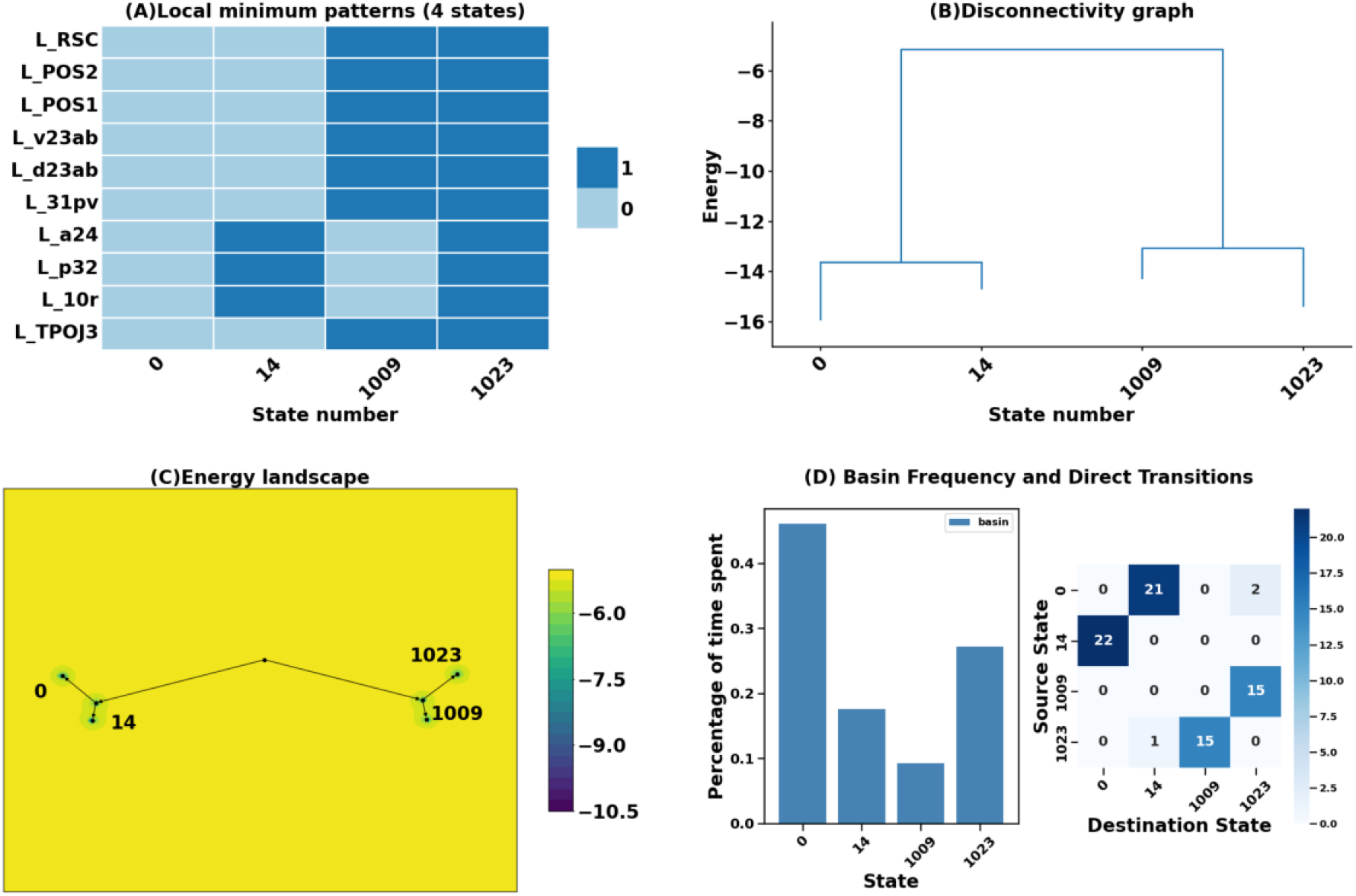
Energy landscape analysis for controls during PCET. (A) Binary activation patterns for 4 local minima across 10 brain regions. (B) Hierarchical disconnectivity graph showing a simpler hierarchical organization with energy values ranging from approximately -6 to -16. (C) Energy landscape visualization showing the spatial organization of local minima and energy gradients across state space. (D) *Left:* Basin frequency distribution showing the percentage of time spent in each basin, with state 0 showing dominant occupancy. *Right:* Direct transition matrix quantifying transition probabilities between states.

### (c) Energy landscape dynamics in AOS

At rest, the AOS energy landscape comprised 7 local minima, 6 of which were observed empirically, with energy values ranging from −6 to −11 **(Figure 3A, B)**, corresponding to fewer, less energetically favorable minima than seen in controls at rest. Occupancy was strongly concentrated: state 0, with all 10 ROIs inactive, was the most frequently visited state at approximately 42% of total time. State 14 accounted for approximately 27% of time, with only 3 of 10 ROIs active (L_a24, L_p32, and L_10r). State 1023 accounted for approximately 15% of time with all 10 ROIs active, and state 497 accounted for approximately 8% with 6 of 10 ROIs active (L_POS2, L_POS1, L_v23ab, L_d23ab, L_31pv, and L_TPOJ3), while the remaining states each falling below a 5% occurrence frequency. The 2D landscape **(Figure 3C)** reflected the relative simplicity of the AOS resting configuration, with 7 nodes organized across a moderately contoured surface with two distinct energy wells corresponding to states 0 and 14. The transition matrix **(Figure 3D)** suggested that state 14 acted as an organizing center, with high numbers of transitions in both directions between states 0, 63, 70, and 1023. State 497 was not directly part of these exchanges and was reached via transitions through state 1023. Notably, the transitions out of state 1023 contrasted with its absorbing status in rest conditions in the control case.

**Figure 3:**
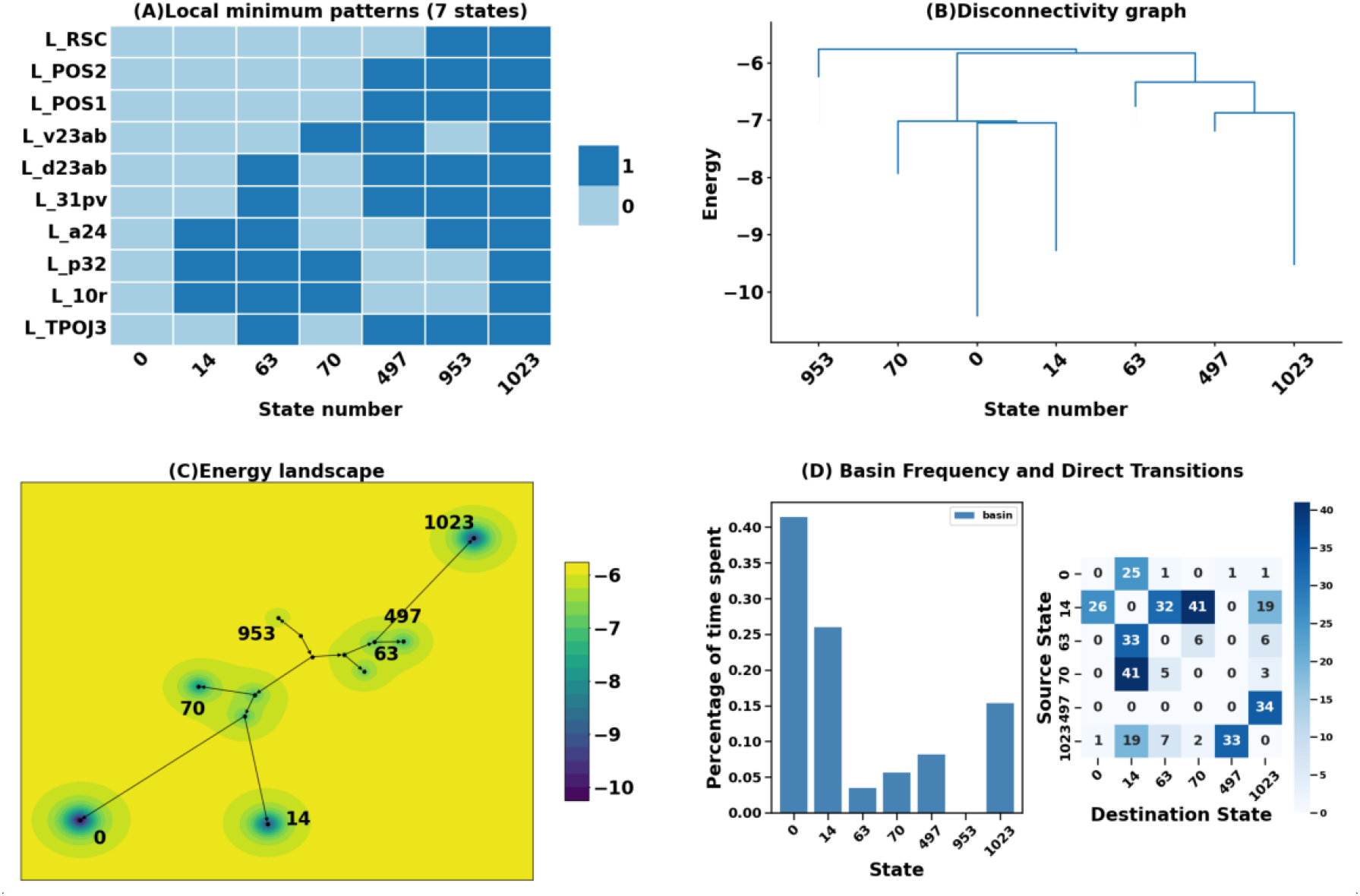
Energy landscape analysis in the resting state for patients. (A) Binary activation patterns for 7 local minima across 10 brain regions. (B) Hierarchical disconnectivity graph showing energy values ranging from -6 to -11. (C) Energy landscape visualization showing the spatial organization of local minima and energy gradients across state space. (D) Left: Basin frequency distribution showing the percentage of time spent in each basin, with states 0, 14, 1023, and 497 most frequently occupied. State 953 was a model-predicted local minimum with zero empirical occupancy. Right: Direct transition matrix quantifying transition probabilities between states.

During PCET, the AOS energy landscape showed a marked expansion rather than consolidation. The number of local minima more than doubled to 15 (Figure 4A, B), with 10 of these states observed empirically and with energy values ranging from −7 to −14, a 114% increase in accessible states in contrast to the 69% decrease seen in controls. Despite this proliferation, actual occupancy remained concentrated in three dominant states: state 0 at approximately 40%, state 14 at approximately 20%, and state 1023 at approximately 15%, with the remaining 12 states collectively accounting for approximately 25% of task time, each individually contributing less than 5%. The dominant state, 0, showed complete DMN suppression with all 10 ROIs inactive. The second most frequent state, 14, was characterized by selective activation of only the anterior cingulate and medial prefrontal regions, with 3 of 10 ROIs active (L_a24, L_p32, and L_10r; all posterior cingulate and temporoparietal regions in the off state). The third most frequent state, 1023, showed complete DMN engagement with all 10 ROIs active (∼15%). Despite its substantial occupancy, state 1023 participated in virtually no direct transitions, suggesting extremely long dwell times once entered—consistent with a deeply trapping attractor state during task performance in AOS. The transition matrix **(Figure 4D)** confirms strong bidirectional coupling between states 0 and 14. Although this matrix contained the highest number of non-zero off-diagonal entries of any condition, all individual transition probabilities remained small, reflecting only occasional excursions away from states 0 and 14 to other states with brief dwell times. Indeed, the main distinction in control task data relative to AOS task data is the consolidation into only 4 accessible states in controls compared to 15 in AOS, with controls showing a stereotyped alternation between the all-off and all-on states, while AOS showed a more dominant inactivation with diffuse transitions across many low-probability configurations.

**Figure 4:**
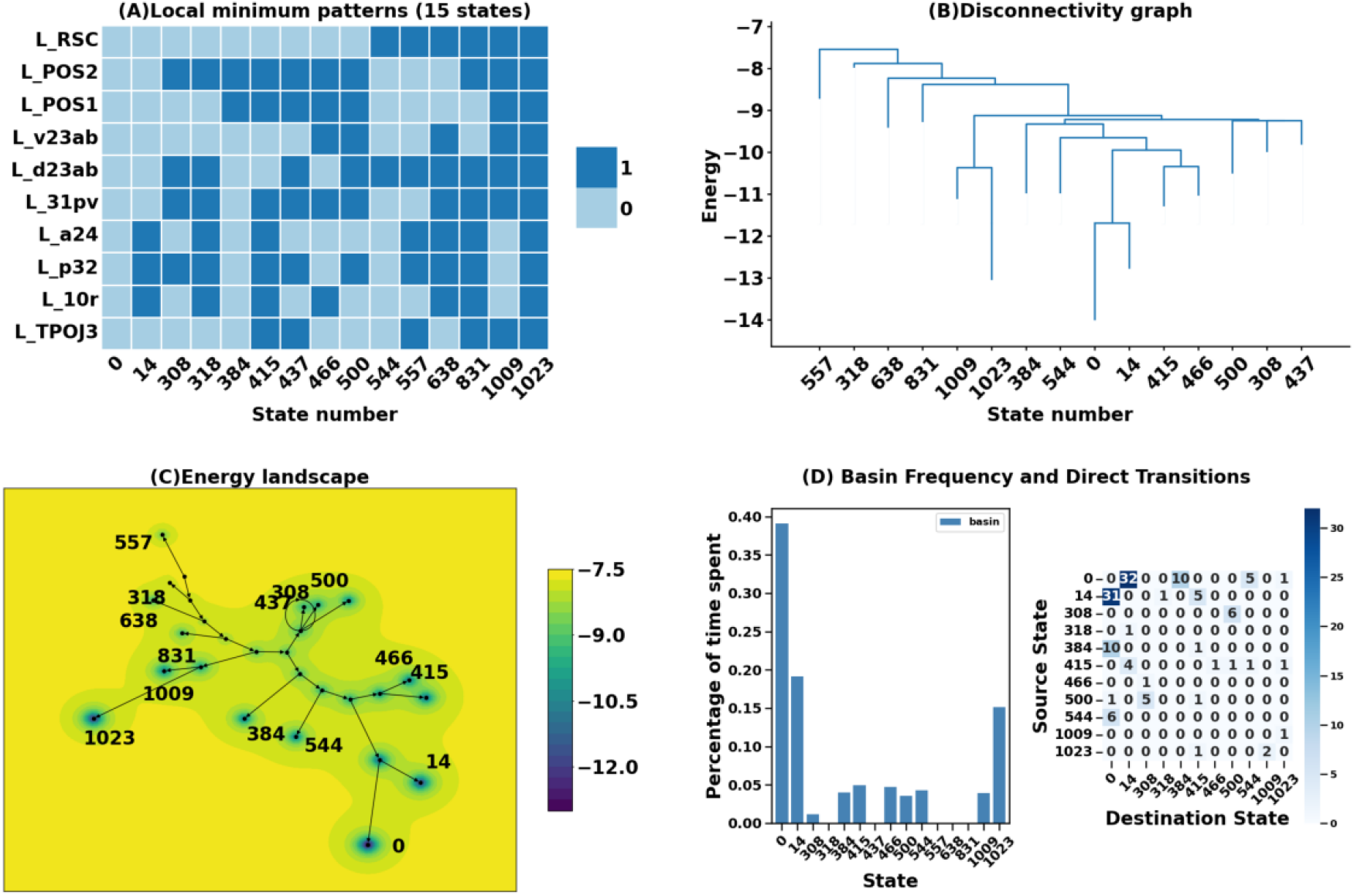
Energy landscape analysis for PCET in patients. (A) Binary activation patterns for 15 local minima across 10 brain regions. (B) Hierarchical disconnectivity graph showing energy barriers between states, with energy values ranging from approximately –7 to –14. (C) Energy landscape visualization showing the spatial organization of local minima and energy gradients across state space. (D) Left: Basin frequency distribution showing the percentage of time spent in each basin, with state 0 (all-off) dominant (∼40%) and state 14 (L_a24, L_p32, L_10r) second most frequent (∼20%). States without bars evident represent model-predicted local minima with zero empirical occupancy. Right: shows direct transition matrix quantifying transition probabilities between minima.

### (d) Between-group comparison and permutation validation

Comparing across groups and conditions reveals interesting patterns of state dominance and transitions. In all cases, landscapes exhibited a dominance of the fully inactive state 0 and state 14 (L_a24, L_p32, and L_10r active) with transitions between the two, as well as frequent appearance of the fully active state 1023. The AOS rest condition landscape was unique, representing the only case with frequent direct transitions between 1023 and the 0/14 complex (specifically, state 14). All four landscapes also included a set of bidirectional transitions between specific dominant states and corresponding less-frequent states, although which states arose differed across populations and conditions.

To confirm that these group differences were not attributable to ROI selection, we conducted permutation tests resampling 500 random sets of 10 ROIs within the DMN and refitting the model each time. One-sample t-tests showed that the observed values for basin number difference and total transitions fell significantly outside the null distributions under both conditions (all p<0.05; **Supplementary Figures 2 and 3**), confirming that the reported landscape differences reflect genuine group-level neural dynamics rather than ROI sampling variability.

## 4. Discussion

Our goal in this study was to conduct ELA of the DMN to gain a more refined understanding of differences between control and AOS populations in dynamic changes in network structure when subjects shift from resting state to perform an attention-demanding task. We observed a markedly different dynamical structure at rest and reorganization of network configuration states when patients and controls switch from resting state to performing an executive function task. At rest, while controls showed a complex energy landscape with a higher number of basins, the AOS showed a simpler landscape with relatively fewer basins. After switching to the task, controls showed a relatively simpler landscape with fewer basins, while AOS showed a dramatically more complex landscape. This shows a fundamentally different network state organization with different responses triggered by an attention-demanding task between the two groups.

From the dynamical systems perspective, in baseline conditions, the DMN has been proposed as a low-energy, high probability attractor state in the brain’s energy landscape (Deco, Jirsa, & McIntosh, 2011; Watanabe, Hirose, et al., 2014). Our analysis also revealed a frequent occurrence of the fully active DMN state (state 1023) during rest periods but added further nuance to our understanding of DMN activation patterns. Indeed, we observed that the most dominant state of the DMN landscape in all conditions (state 0) featured inactivity in all ROIs. Moreover, the relative frequency of the three dominant states, all-on (1023), all-off (0), and a third state (14) with activity in three anterior cingulate and medial prefrontal regions (L_a24, L_p32, and L_10r), did not change between rest and task conditions in either population. However, the all-on state was relatively more frequent in controls than in patients and in rest conditions; it acted as a nearly-absorbing state, consistent with its previous characterization as a baseline attractor. Thus, our study highlights first that the DMN is not simply a static network active at rest and suppressed during tasks and second that AOS patients may have difficulty sustaining fully active DMN states relative to controls.

Distinct RSNs have been proposed to be organized with different dynamical principles, with DMN found to be organized around flexibility, while others such as frontoparietal networks favor stability (Watanabe, Hirose, et al., 2014). According to these principles, efficient cognition depends on the ability of the DMN to transition out of its baseline attracting state into task-relevant configurations, which correspond to suppression of DMN in healthy individuals that has been documented as in schizophrenia (Menon, 2023; Whitfield-Gabrieli & Ford, 2012). In our analysis, alterations in the control of the DMN manifest at the landscape level not as the absence of an all-off state, but as a failure to consolidate task engagement into a small set of stable, organized configurations. Both groups retained the all-off state as the dominant configuration during PCET, but controls achieved this through a dramatically simplified landscape of only 4 states with stereotyped alternation between complete suppression and complete engagement that together accounted for ∼74% of PCET time, with the all-off state occupying the energetically deepest minimum (Figure 2). This favoring of the all-off state is consistent with the coordinated deactivation required to redirect resources toward task-relevant networks (Cole et al., 2014). Thus, the finding of suppression of DMN regions during task engagement is overly simplistic when drawn from traditional functional connectivity and regional BOLD response analyses, and our findings about transitions between states highlight the underlying dynamics of the suppression.

In AOS, by contrast, while the all-off state remained the most frequently visited configuration during PCET (∼40%), it was accompanied by 14 additional accessible configurations, with the second most frequent state being one of selective anterior cingulate and medial prefrontal activation (L_a24, L_p32, and L_10r). The all-on state was also frequently observed; in contrast to controls during PCET, however, this fully active state participated in very few transitions. Task performance in AOS therefore appears to trap the system in the fully active DMN configuration for prolonged periods, which may reflect an inability to flexibly disengage from full DMN activation during cognitive demand. Such observations can only be discovered in a state-space model. Prior ELA studies in schizophrenia and mood disorders have reported altered landscape structure (Ishida et al., 2024), but the present data identify the specific failure of adaptation when patients shift from rest to task engagement with an inability to consolidate into a small number of stable task-relevant configurations and to adjust DMN activation to achieve functional states.

Our finding that the DMN in AOS maintains a relatively consolidated organization, but with a disorganization in network reconfiguration for executive task performance, aligns with previous computational work. Specifically, computational models of schizophrenia have proposed that cognitive deficits arise from disrupted temporal organization of brain states rather than from altered connection strengths per se (Deco et al., 2015; Rolls & Deco, 2011). Our findings provide direct empirical support for this idea by showing the divergence of the number and character of accessible states between groups at rest and under task conditions. These results are consistent with prior ELA work in this dataset showing that AOS trajectories traverse higher-energy states during executive function tasks (Theis et al., 2025) and extend that finding by identifying the landscape-level mechanism of proliferation of shallow local minima rather than global elevation of energy in AOS during task performance.

The transition matrices that we derived provide further insights beyond those obtained from counting frequently observed states and quantifying their occupancy rates. We have already discussed the lack of transitions out of the fully active state in the AOS task condition. In controls, during PCET, transitions were concentrated into two dominant pathways, indicating that the system followed a small number of stereotyped, high-probability sequences between configurations, a signature of reliable, flexible, and repeatable network dynamics. In AOS during PCET, transitions were primarily concentrated between two dominant states but with a diffuse distribution of additional, low-probability transitions across many state pairs. This loss of stereotyped transition pathways is mechanistically important because cognitive performance depends not only on occupying appropriate states but also in transitioning between them in a reliable manner (Uhlhaas & Singer, 2012). The anomalous excursions visible in the AOS transition matrix associated with task performance (Figure 4D) reflect the ∼25% of PCET time distributed across 12 low-probability configurations, each arising during less than 5% of the observation time, suggesting that the system frequently departed from its two dominant states into a wide variety of transient configurations rather than following stereotyped transition pathways, a pattern unlikely to support consistent executive function performance. These findings suggest that interventions targeting dynamical instability specifically under cognitive load may be more effective than those aimed at resting-state connectivity. Our prior work showed that cognitive enhancement therapy can favorably alter pairwise functional connectivity in schizophrenia (Keshavan, Eack, Prasad, Haller, & Cho, 2017) despite evidence of reduced structural network resilience in this population (Lewis et al., 2023). The present framework suggests that task-state landscape consolidation, specifically the emergence of a stable DMN suppression attractor and concentrated transition pathways during PCET, could serve as a more mechanistically grounded outcome measure for such interventions.

Strengths of our study include acquisition of imaging data at ultra-high field strength (7T), providing superior signal-to-noise ratio compared to lower field magnets. This could enable more precise characterization of DMN activity patterns. Second, both resting-state and task fMRI were acquired from the same individuals within the same scanning session, ensuring that the crossover interaction between groups and conditions cannot be attributed to cohort differences or inter-session variability. Third, we have examined both resting and task fMRI data together to elucidate the dynamical features of the energy landscape. Fourth, we have examined a relatively severe form of schizophrenia that may feature a greater impact of neurodevelopmental mechanisms (Kuo, Musket, et al., 2022; Kuo, Roalf, et al., 2022). Finally, the robustness of the energy landscape findings was confirmed through permutation testing, demonstrating that the reported group differences are not attributable to ROI selection bias.

Some limitations are that the ELA was performed at the group level by concatenating time series across subjects, because individual scan lengths provide insufficient coverage of the 1,024-dimensional binary state space, reflecting the exponential scaling of state space with network size, to allow for reliable individual-level parameter estimation, as was observed by others (Ezaki et al., 2017; Ishida et al., 2024; Watanabe & Rees, 2017). We did not correlate energy levels and transition dynamics since we had previously published on the association of higher energy with poorer executive functions when executive function network was examined (Theis et al., 2025). Future work employing Bayesian approaches to individual-level estimation or longer scanning protocols may enable subject-level analyses and formal hypothesis testing. The cross-sectional design additionally prevents examination of whether landscape organization normalizes with treatment or tracks disease progression, questions that longitudinal application of this framework could address.

## Supplemental Materials

**Figure 1.**
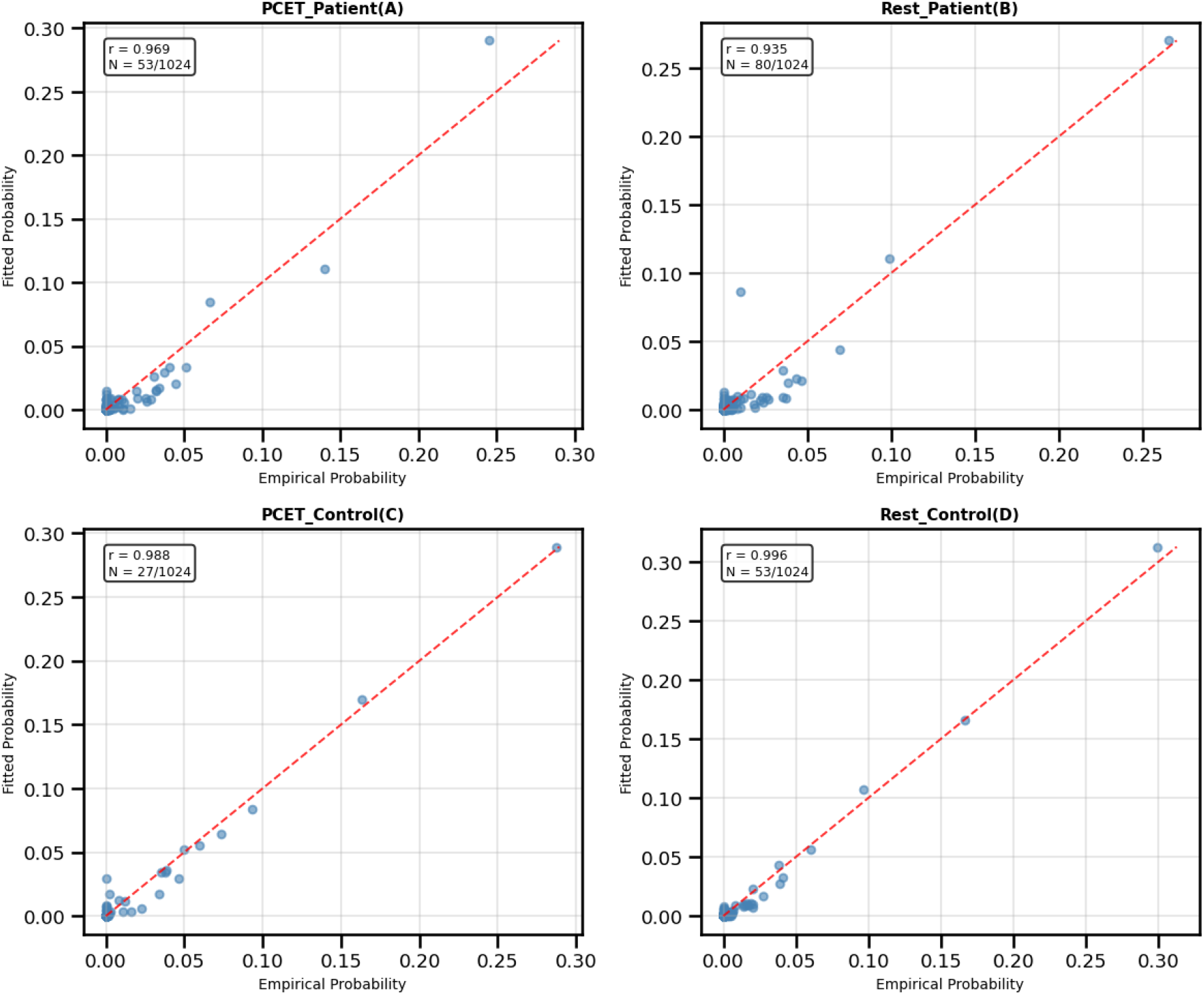
Model fit quality for PCET and Resting-state data. Empirical vs. fitted state probabilities (A) for PCET patients (*r* = 0.969, 53/1024 states observed), (C) healthy controls (*r* = 0.988, 27/1024 states observed), (B) Resting-state patients (*r* = 0.935, 80/1024 states observed) and (D) Resting-state healthy controls (*r*=0.996, 53/1024 states observed). Red dashed line indicates perfect fit. Note: *N* denotes the number of unique brain states observed out of 2^10^ possible configurations.

**Figure 2.**
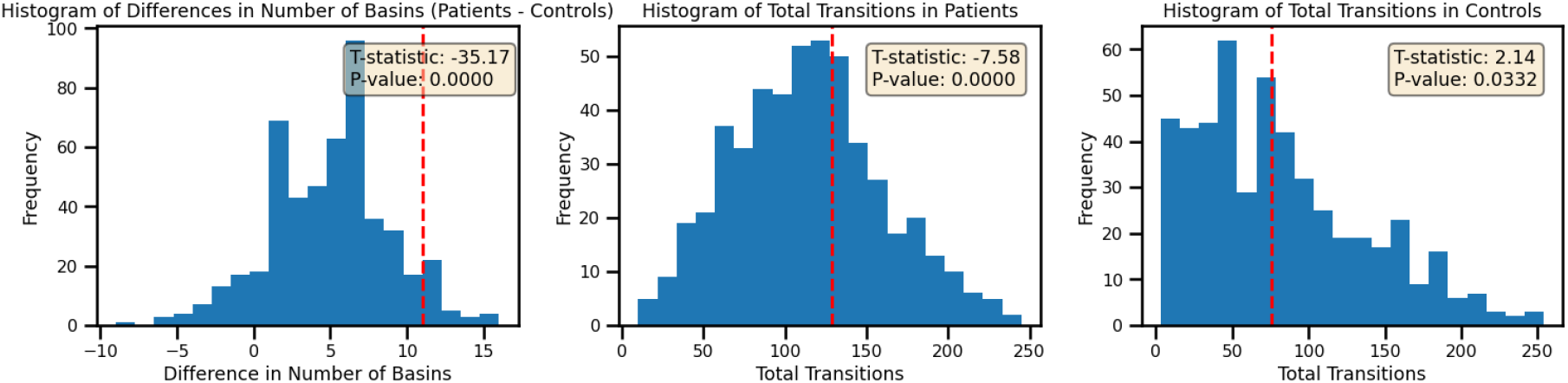
Permutation test results for the PCET condition. Null distributions (blue histograms) were generated by resampling 10 ROIs from patient and control datasets across 500 iterations, with Ising models fitted to each sample. Red dashed lines indicate the observed values from the full analysis. Left: distribution of patient-minus-control differences in number of energy basins (observed = 11; t = -35.17, p < 0.0001). Middle: distribution of total state transitions in patients (observed = 129; t = -7.58, p < 0.0001). Right: distribution of total state transitions in controls (observed = 76; t = 2.14, p = 0.033). Observed values falling outside the null distributions confirm that group differences in energy landscape topology are not attributable to ROI sampling variability.

**Figure 3.**
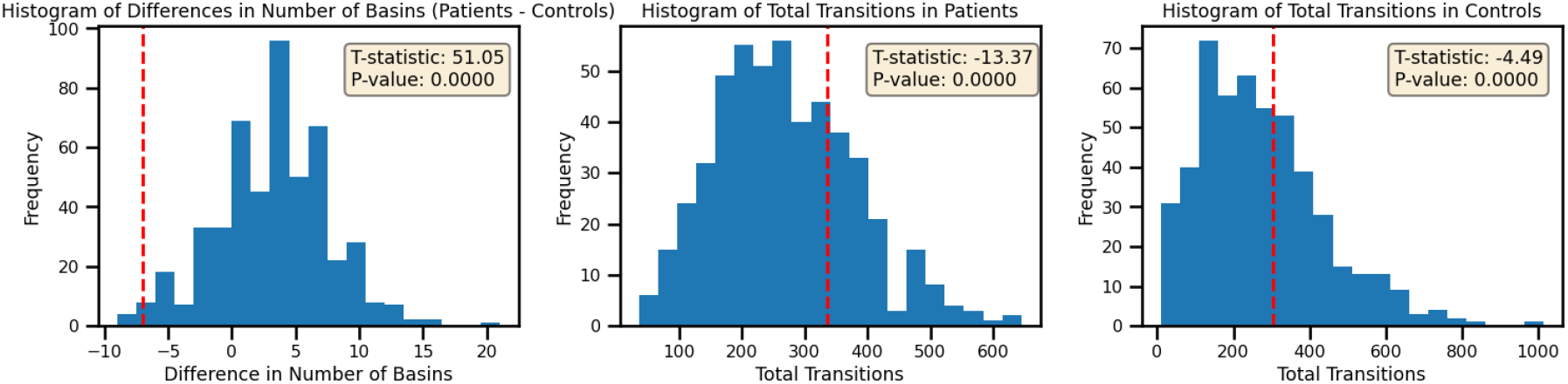
Permutation test results for the resting state condition. Null distributions (blue histograms) were generated using the same resampling procedure as in the PCET condition. Red dashed lines indicate the observed values from the full analysis. Left: distribution of patient-minus-control differences in number of energy basins (observed = −7; t = 51.05, p < 0.0001). Middle: distribution of total state transitions in patients (observed = 320; t = −13.37, p < 0.0001). Right: distribution of total state transitions in controls (observed = 350; t = −4.49, p < 0.0001). All observed values deviate significantly from their respective null distributions, supporting the robustness of the resting-state energy landscape findings.

## REFERENCES

Brickman, A. M., Buchsbaum, M. S., Bloom, R., Bokhoven, P., Paul-Odouard, R., Haznedar, M. M., … Shihabuddin, L. (2004). Neuropsychological functioning in first-break, never-medicated adolescents with psychosis. J Nerv Ment Dis, 192(9), 615–622.

Brush, S. G. (1967). History of the Lenz-Ising Model. Reviews of Modern Physics, 39(4), 883–893. doi:10.1103/RevModPhys.39.883

Cole, Michael W., Bassett, Danielle S., Power, Jonathan D., Braver, Todd S., & Petersen, Steven E. (2014). Intrinsic and Task-Evoked Network Architectures of the Human Brain. Neuron, 83(1), 238–251. doi:10.1016/j.neuron.2014.05.014

Deco, G., Jirsa, V. K., & McIntosh, A. R. (2011). Emerging concepts for the dynamical organization of resting-state activity in the brain. Nat Rev Neurosci, 12(1), 43–56. doi:10.1038/nrn2961

Deco, G., Tononi, G., Boly, M., & Kringelbach, M. L. (2015). Rethinking segregation and integration: contributions of whole-brain modelling. Nat Rev Neurosci, 16(7), 430–439. doi:10.1038/nrn3963

Dijkstra, E. W. (1959). A note on two problems in connexion with graphs. Numerische Mathematik, 1(1), 269–271. doi:10.1007/BF01386390

Esteban, O., Markiewicz, C. J., Blair, R. W., Moodie, C. A., Isik, A. I., Erramuzpe, A., … Gorgolewski, K. J. (2019). fMRIPrep: a robust preprocessing pipeline for functional MRI. Nat Methods, 16(1), 111–116. doi:10.1038/s41592-018-0235-4

Ezaki, T., Watanabe, T., Ohzeki, M., & Masuda, N. (2017). Energy landscape analysis of neuroimaging data. Philos Trans A Math Phys Eng Sci, 375(2096), 20160287. doi:10.1098/rsta.2016.0287

Fischl, B. (2012). FreeSurfer. Neuroimage, 62(2), 774–781. doi:10.1016/j.neuroimage.2012.01.021

Frangou, S. (2010). Cognitive function in early onset schizophrenia: a selective review. Front Hum Neurosci, 3, 79. doi:10.3389/neuro.09.079.2009

Friston, K. J., Harrison, L., & Penny, W. (2003). Dynamic causal modelling. Neuroimage, 19(4), 1273–1302. doi:10.1016/s1053-8119(03)00202-7

Glasser, M. F., Coalson, T. S., Robinson, E. C., Hacker, C. D., Harwell, J., Yacoub, E., … Van Essen, D. C. (2016). A multi-modal parcellation of human cerebral cortex. Nature, 536(7615), 171–178. doi:10.1038/nature18933

Glasser, M. F., Sotiropoulos, S. N., Wilson, J. A., Coalson, T. S., Fischl, B., Andersson, J. L., … Jenkinson, M. (2013). The minimal preprocessing pipelines for the Human Connectome Project. Neuroimage, 80, 105–124. doi:10.1016/j.neuroimage.2013.04.127

Gur, R. C., Ragland, J. D., Moberg, P. J., Bilker, W. B., Kohler, C., Siegel, S. J., & Gur, R. E. (2001). Computerized neurocognitive scanning: II. The profile of schizophrenia. Neuropsychopharmacology, 25(5), 777–788.

Gur, R. C., Ragland, J. D., Moberg, P. J., Turner, T. H., Bilker, W. B., Kohler, C., … Gur, R. E. (2001). Computerized neurocognitive scanning: I. Methodology and validation in healthy people. Neuropsychopharmacology, 25(5), 766–776.

Holtmaat, A., & Svoboda, K. (2009). Experience-dependent structural synaptic plasticity in the mammalian brain. Nat Rev Neurosci, 10(9), 647–658. doi:10.1038/nrn2699

Ishida, T., Yamada, S., Yasuda, K., Uenishi, S., Tamaki, A., Tabata, M., … Kimoto, S. (2024). Aberrant brain dynamics of large-scale functional networks across schizophrenia and mood disorder. NeuroImage: Clinical, 41, 103574. doi:10.1016/j.nicl.2024.103574

Ising, E. (1924). Beitrag zur Theorie des Ferro- und Paramagnetismus. University of Hamburg, Retrieved from https://www.icmp.lviv.ua/ising/books/isingshort.pdf

Ising, T., Folk, R., Kenna, R., Berche, B., & Holovatch, Y. (2017). The Fate of Ernst Ising and the Fate of his Model. arXiv: History and Philosophy of Physics.

Karatekin, C., Bingham, C., & White, T. (2009). Regulation of cognitive resources during an n-back task in youth-onset psychosis and attention-deficit/hyperactivity disorder (ADHD). Int J Psychophysiol, 73(3), 294–307. doi:10.1016/j.ijpsycho.2009.05.001

Karatekin, C., White, T., & Bingham, C. (2008). Divided attention in youth-onset psychosis and attention deficit/hyperactivity disorder. J Abnorm Psychol, 117(4), 881–895. doi:10.1037/a0013446

Kaufman, J., Birmaher, B., Brent, D. A., Ryan, N. D., & Rao, U. (2000). K-Sads-Pl. J Am Acad Child Adolesc Psychiatry, 39(10), 1208.

Keshavan, M. S., Eack, S. M., Prasad, K. M., Haller, C. S., & Cho, R. Y. (2017). Longitudinal functional brain imaging study in early course schizophrenia before and after cognitive enhancement therapy. Neuroimage, 151, 55–64. doi:10.1016/j.neuroimage.2016.11.060

Kester, H. M., Sevy, S., Yechiam, E., Burdick, K. E., Cervellione, K. L., & Kumra, S. (2006). Decisionmaking impairments in adolescents with early-onset schizophrenia. Schizophr Res, 85(1-3), 113–123. doi:S0920-9964(06)00102-2 [pii]; 10.1016/j.schres.2006.02.028

Kumra, S., & Charles Schulz, S. (2008). Editorial: research progress in early-onset schizophrenia. Schizophr Bull, 34(1), 15–17. doi:sbm123 [pii]; 10.1093/schbul/sbm123

Kumra, S., Shaw, M., Merka, P., Nakayama, E., & Augustin, R. (2001). Childhood-onset schizophrenia: research update. Can J Psychiatry, 46(10), 923–930.

Kuo, S. S., Musket, C. W., Rupert, P. E., Almasy, L., Gur, R. C., Prasad, K. M., … Pogue-Geile, M. F. (2022). Age-dependent patterns of schizophrenia genetic risk affect cognition. Schizophr Res, 246, 39–48. doi:10.1016/j.schres.2022.05.012

Kuo, S. S., Roalf, D. R., Prasad, K. M., Musket, C. W., Rupert, P. E., Wood, J., … Pogue-Geile, M. F. (2022). Age-dependent effects of schizophrenia genetic risk on cortical thickness and cortical surface area: Evaluating evidence for neurodevelopmental and neurodegenerative models of schizophrenia. J Psychopathol Clin Sci, 131(6), 674–688. doi:10.1037/abn0000765

Lewis, M., Santini, T., Theis, N., Muldoon, B., Dash, K., Rubin, J., … Prasad, K. (2023). Modular architecture and resilience of structural covariance networks in first-episode antipsychotic-naive psychoses. Sci Rep, 13(1), 7751. doi:10.1038/s41598-023-34210-y

Menon, V. (2023). 20 years of the default mode network: A review and synthesis. Neuron, 111(16), 2469–2487. doi:10.1016/j.neuron.2023.04.023

Musket, C. W., Kuo, S. S., Rupert, P. E., Almasy, L., Gur, R. C., Prasad, K., … Pogue-Geile, M. F. (2020). Why does age of onset predict clinical severity in schizophrenia? A multiplex extended pedigree study. Am J Med Genet B Neuropsychiatr Genet, 183(7), 403–411. doi:10.1002/ajmg.b.32814

Petersen, A. C., Crockett, L., Richards, M., & Boxer, A. (1988). A Self-Report Measure of Pubertal Status: Reliability, Validity, and Initial Norms. Journal of Youth and Adolescence, 17(2), 117–133.

Rapoport, J. L., Giedd, J., Kumra, S., Jacobsen, L., Smith, A., Lee, P., … Hamburger, S. (1997). Childhood-onset schizophrenia. Progresive ventricular change during adolescence. Arch Gen Psychiatry, 54(10), 897–903.

Rapoport, J. L., & Gogtay, N. (2011). Childhood onset schizophrenia: support for a progressive neurodevelopmental disorder. Int J Dev Neurosci, 29(3), 251–258. doi:10.1016/j.ijdevneu.2010.10.003

Rolls, E. T., & Deco, G. (2011). A computational neuroscience approach to schizophrenia and its onset. Neurosci Biobehav Rev, 35(8), 1644–1653. doi:10.1016/j.neubiorev.2010.09.001

Sandhu, Z., Tanglay, O., Young, I. M., Briggs, R. G., Bai, M. Y., Larsen, M. L., … Sughrue, M. E. (2021). Parcellation-based anatomic modeling of the default mode network. Brain Behav, 11(2), e01976. doi:10.1002/brb3.1976

Tang, A., Jackson, D., Hobbs, J., Chen, W., Smith, J. L., Patel, H., … Beggs, J. M. (2008). A maximum entropy model applied to spatial and temporal correlations from cortical networks in vitro. J Neurosci, 28(2), 505–518. doi:10.1523/JNEUROSCI.3359-07.2008

Thaden, E., Rhinewine, J. P., Lencz, T., Kester, H., Cervellione, K. L., Henderson, I., … Kumra, S. (2006). Early-onset schizophrenia is associated with impaired adolescent development of attentional capacity using the identical pairs continuous performance test. Schizophr Res, 81(2-3), 157–166. doi:10.1016/j.schres.2005.09.015

Theis, N., Bahuguna, J., Rubin, J. E., Banerjee, S. S., Muldoon, B., & Prasad, K. M. (2025). Energy of Functional Brain States Correlates With Cognition in Adolescent-Onset Schizophrenia and Healthy Persons. Hum Brain Mapp, 46(1), e70129. doi:10.1002/hbm.70129

Uhlhaas, P. J., & Singer, W. (2012). Neuronal dynamics and neuropsychiatric disorders: toward a translational paradigm for dysfunctional large-scale networks. Neuron, 75(6), 963–980. doi:10.1016/j.neuron.2012.09.004

Watanabe, T., Hirose, S., Wada, H., Imai, Y., Machida, T., Shirouzu, I., … Masuda, N. (2014). Energy landscapes of resting-state brain networks. Front Neuroinform, 8, 12. doi:10.3389/fninf.2014.00012

Watanabe, T., Masuda, N., Megumi, F., Kanai, R., & Rees, G. (2014). Energy landscape and dynamics of brain activity during human bistable perception. Nat Commun, 5, 4765. doi:10.1038/ncomms5765

Watanabe, T., & Rees, G. (2017). Brain network dynamics in high-functioning individuals with autism. Nat Commun, 8, 16048. doi:10.1038/ncomms16048

White, T., Mous, S., & Karatekin, C. (2013). Memory-guided saccades in youth-onset psychosis and attention deficit hyperactivity disorder (ADHD). Early Interv Psychiatry. doi:10.1111/eip.12038

Whitfield-Gabrieli, S., & Ford, J. M. (2012). Default mode network activity and connectivity in psychopathology. Annu Rev Clin Psychol, 8, 49–76. doi:10.1146/annurev-clinpsy-032511-143049

